# Moistube™ Irrigation (MTI) Discharge Under Variable Evaporative Demand

**DOI:** 10.1101/2020.07.02.184309

**Authors:** Tinashe Lindel Dirwai, Aidan. Senzanje, Tafadzwanashe. Mabhaudhi

**Author notes:** Corresponding author (TLD).

## Abstract

We investigated the conceptual capability of Moistube irrigation to discharge under zero applied positive pressure under varied climatic conditions by inducing an artificial evaporative demand (*E*_*d*_) or negative pressure around moistube tubing. This was premised on the null hypothesis that an artificially induced *E*_*d*_ or negative pressure does not impact moistube discharge. Moistube tubing was enclosed in a 1 m long PVC conduit. A 20 l water reservoir placed on an electronic balance provided a continuous supply of water whilst a three-speed hot air blower facilitated the radiative factor and advection process. The procedure was conducted under varied climatic conditions of air velocities (*u*_*a*_) 1.2 m.s^−1^, 2.5 m.s^−1^, and 3.0 m.s^−1^ and the experiment run times were 159 h, 134 h and 10 h, respectively. The average temperature (*T*_*ave*_) and relative humidity (RH) data for *u*_*a*_ = 1.2 m.s^−1^ were 53°C and 7.31%, whilst for *u*_*a*_ = 2.5 m.s^−1^, *T*_*ave*_ was 56°C and RH = 7.19%, and for *u*_*a*_ = 3.0 m.s^−1^, *T*_*ave*_ was 63°C and RH = 6.16%. The experimental data was input into the four variable Penman-Monteith method to compute the evaporative demand (*E*_*d*_). For each of the air velocities, the respective *E*_*d*_ values obtained were 0.16, 0.31 and 0.36 mm.d^−1^. The Bowen ratios (*r*) were well below 1 (*r* < 1), which suggested a sufficient supply of moisture to evaporate. For *E*_*d*_ = 0.16 mm.d^−1^ the vapour pressure deficit (VPD) was 113.08 mbars, whilst for *E*_*d*_ = 0.31 mm.d^−1^ and for *E*_*d*_ = 0.36 mm.d^−1^ the VPD were 129.93 mbars and 150.14 mbars, respectively. The recorded discharges (*q*) at *t* = 10 hrs for *E*_*d*_= 0.16 mm.d^−1^ was 7.67*10^−3^ l.hr^−1^.m^−1^, whilst for *E*_*d*_ = 0.31 mm.d^−1^ *q* = 14.5*10^−3^ l.hr^−1^.m^−1^, and for *E*_*d*_ = 0.36 mm.d^−1^ *q* = 20.8*10^−3^ l.hr^−1^.m^−1^. The imposed negative pressure causes an exponential increase in moistube discharge, thus disproving the null hypothesis. The higher the evaporative demand the higher the discharge. This phenomenon allows moistube irrigation to be used for deficit irrigation purposes and allows irrigators to capitalize on realistic soil matric potential irrigation scheduling approach.

## 1 Introduction

Moistube™ irrigation (MTI) is a relatively new semi-permeable membrane (SPM) irrigation technology. A typical third generation Moistube™ pipe has an outer protective membrane and an inner membrane that constitutes of densely and uniformly spaced nano-pores whose pore-diameter ranges from 10 – 900 nm. The technology utilises nano-technology such that the inner membrane imitates plant water uptake, which facilitates discharge according to crop water requirements (1, 2). It is a low pressure discharge sub-surface irrigation technology whose functionality is similar to ceramic pitcher pots. Under a negative pressure, or in the absence of applied pressure, the discharge is a function of matric potential (*ψ*) (1–3). Conceptually, when water potential (*ψ*_*water*_) is greater than the matric potential of the surrounding soil (*ψ*_*soil*_), the MTI discharge rate is high and vice versa (1, 3). There exists a number of issues in need of research answers, for example, how MTI discharge varies with imposed evaporative demand (*E*_*d*_). The *E*_*d*_ mimics changing soil water conditions thus exploring MTI applicability to deficit irrigation.

According to (<u>4</u>), evapotranspiration (*ET*) is a combination of water loss from soils and transpiration by plants; the water loss mechanisms occur simultaneously under ambient conditions. A simpler method of estimating *ET* is by using evaporative demand (*E*_*d*_), which is defined as the upper boundary for *ET* under ambient conditions with an uncapped hydrological limit or unlimited water supply. *E*_*d*_ is used in irrigation scheduling as a proxy for plant water consumptive use wherein crop coefficients or factors such as phenology and soil stress are used to estimate the *ET*_*o*_ (5). There exists a correlation between rate and amount of plant water use and *E*_*d*_ (6), thus since MTI discharge is a function of soil matric potential a high *E*_*d*_ potentially increase MTI discharge. Currently there is a dearth in literature concerning porous and continuous emitter discharge and *E*_*d*_. What is known is how rate and amount of water uptake correlates with *E*_*d*_.

*E*_*d*_ has three drivers, which are the hydrological, radiative and the advective limits (5, 7). For evaporative demand to occur there has to be an adequate water supply to meet the minimum hydrological limit hence, the hydrological limit defines the availability of water to evaporate and transpire from plants and soil surfaces. <u>Hobbins and Huntington (5)</u> mathematically characterised it as in Equation 1:

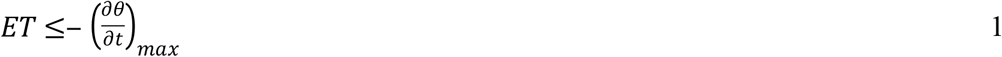

where: *ET* = evapo-transpiration (mm.d^−1^), 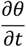 = time rate change of moisture availability in units of mass flux, and *max* = time taken by soil to avail water for evaporation (exfiltrate) (mm.d^−1^).

According to <u>Hobbins and Huntington (5)</u>, the radiative limit is the energy required to facilitate the evaporative process and it is modelled according to Equation 2:

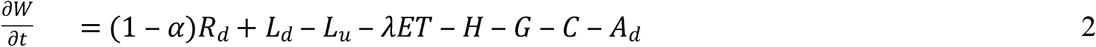

where: 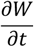 = time rate change of heat (*W*) storage in evaporating layer, α = surface albedo (constant), *R*_*d*_ = downward shortwave radiation incident at the surface, *L*_*u*_ = longwave radiation outward from the surface, *L*_*d*_ = longwave radiation inward to the surface, λET = latent heat flux, *H* = sensible heat flux, *G* = net heat flux conducted from the evaporating surface into the soil, *C* = energy absorbed by vegetation in the control volume, and *A*_*d*_ = heat gained by advection to the control volume.

All terms in Equation 2 are in flux units (W.m^−2^, multiply by 86,400/λ for mm day^−1^ where λ = latent heat of vaporization of water, J kg^−1^) (5). A typical buried MTI lateral tubing and the active heat fluxes is depicted in Figure 1.

**Fig 1.**
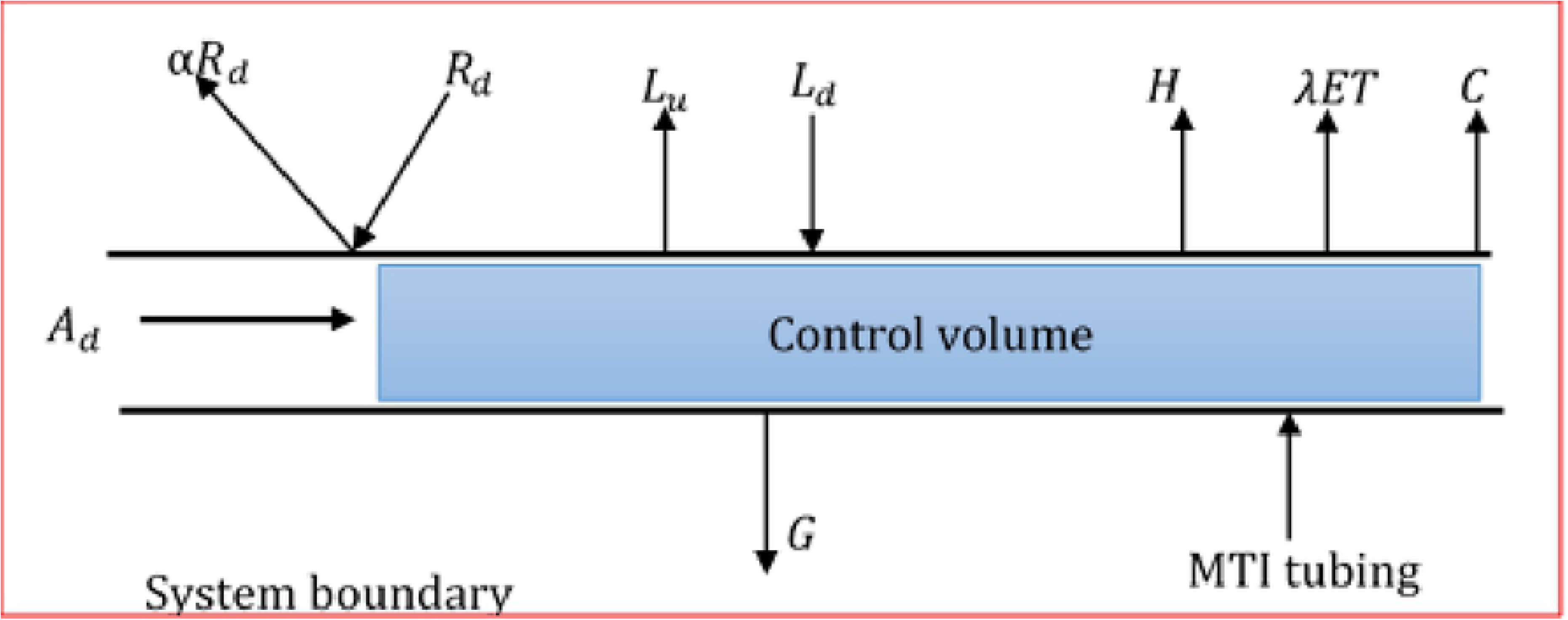
Conceptual active fluxes acting on a MTI lateral buried in the soil.

The advective limit describes the system boundary’s ability to absorb and bear away moisture and it can mathematically be modelled as in Equation 3 (<u>5</u>):

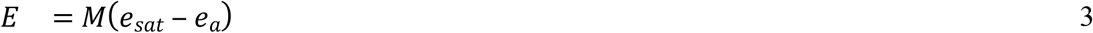

where: *E* = evaporation rate (mm.d^−1^), *e*_*sat*_ = saturation vapour pressure at water temperature (mbars), *e*_*a*_ = saturation vapour pressure of air (mbars), and *M* = mass transfer coefficient, which can be expressed as a function of air velocity (*u*_*a*_), which dictates the variable vapour pressure as in Equation 4.

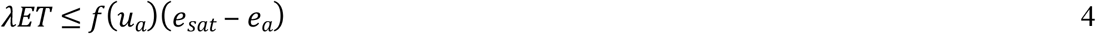

where *f*(*u*_*a*_) is measured in (W.m-^2^.N.m^−2^.).

Limited research efforts have been made to model *E*_*d*_ under controlled and varied micro-climatic conditions. For example, <u>Donohue, McVicar (7)</u> used five evapotranspiration formulations namely Penman, Priestley–Taylor, Morton point, Morton areal and Thornthwaite to model and assess the best proxy for *E*_*d*_.

<u>Abu-Zreig, Zraiqat (8)</u> modelled an artificially induced pan evaporation scenario in order to measure the discharge rates of ceramic pitchers under negative head. The theoretical discharge design aspects of the MTI technology have not been tested. Understanding negative pressure discharge capability of MTI tubing can potentially aid irrigators to inform deficit irrigation strategies and save on energy costs that otherwise drive positive head irrigation systems.

The study investigated the conceptual design and supposed discharge mechanism of Moistube™ when subjected to a negative pressure or in the absence of a positive driving pressure. The study hypothesized that imposed negative pressure or an artificial *E*_*d*_ cannot induce MTI discharge. This study adds to knowledge by providing answers around the conceptual zero pressure head discharge capability of MTI. The *E*_*d*_ was used to simulate a buried Moistube™ tubing under low matric potential or negative soil water tension conditions whilst monitoring the subsequent discharge performance.

## 2 Materials and Methods

### 2.1 Study Site

The experiment was carried out at the University of KwaZulu-Natal Hydrology Laboratory (29.626044, 30.403325). The laboratory had a controlled room temperature of 22°C ± 1°C and a measured relative humidity (RH) of 55% ± 5%. Controlled conditions helped to eliminate the variations in atmospheric temperature and humidity.

### 2.2 Experimental Design and Set-up

#### 2.2.1 Design

The experiment was a single factor experiment (*E*_*d*_). The experiment constituted of five recorded variables, namely, relative humidity (RH), air velocity (*u*_*a*_), net radiation (*R*_*n*_), mass flow rate (*m*), and micro-climate temperature (*T*_*a*_). The experiment had three controlled *u*_*a*_ velocities; 1.2 m.s^−1^, 2.5 m.s^−1^ and 3.0 m.s^−1^. The *u*_*a*_ selection was dictated by the default hot air blower settings. For each *u*_*a*_, the subsequent *T*_*a*_, *R*_*n*_, and vapour pressure deficit (VPD) derived from RH and *T*_*a*_ was recorded at five-minute intervals for 159 hr, 134 hr and 10 hr. Thereafter, the resultant *E*_*d*_ was computed. The VPD was expressed as the difference between actual vapour pressure of the air (*e*_*a*_) and the observed vapour pressure (*e*). The *m* was converted to discharge (*q*) by multiplying the recorded values by the density of water (*p*_*w*_). The experimental variables are shown in Figure 2 and setup in Figure 3.

**Fig 2.**
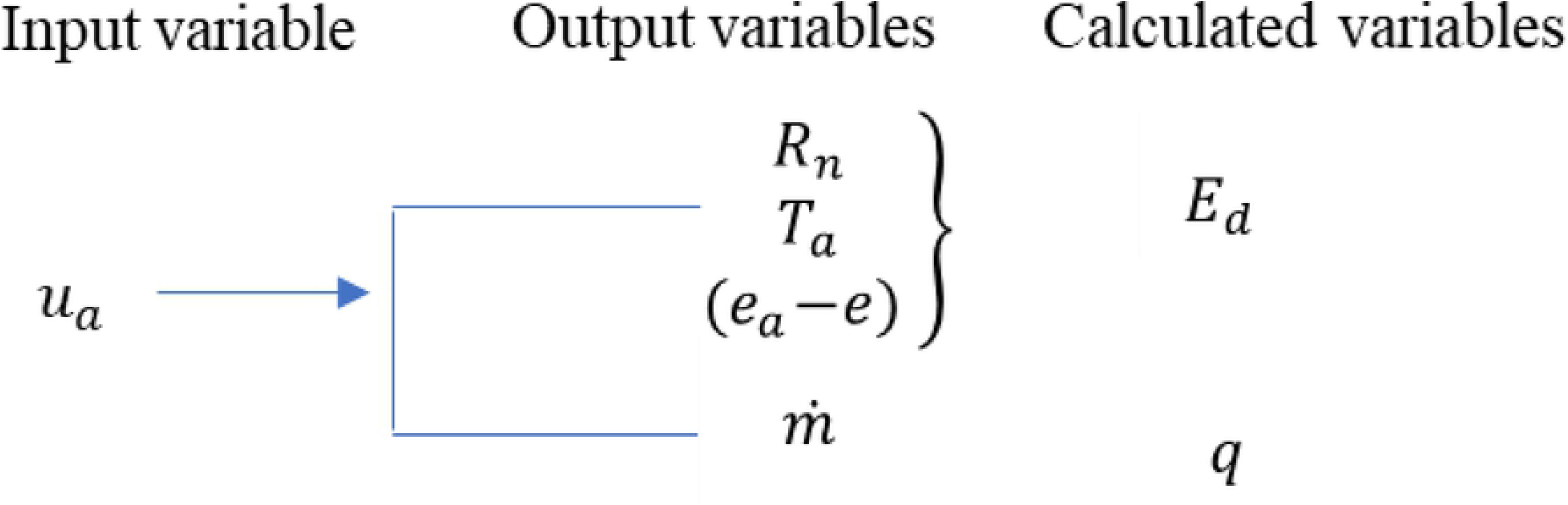
The recorded evaporative demand experimental parameters

**Fig 3.**
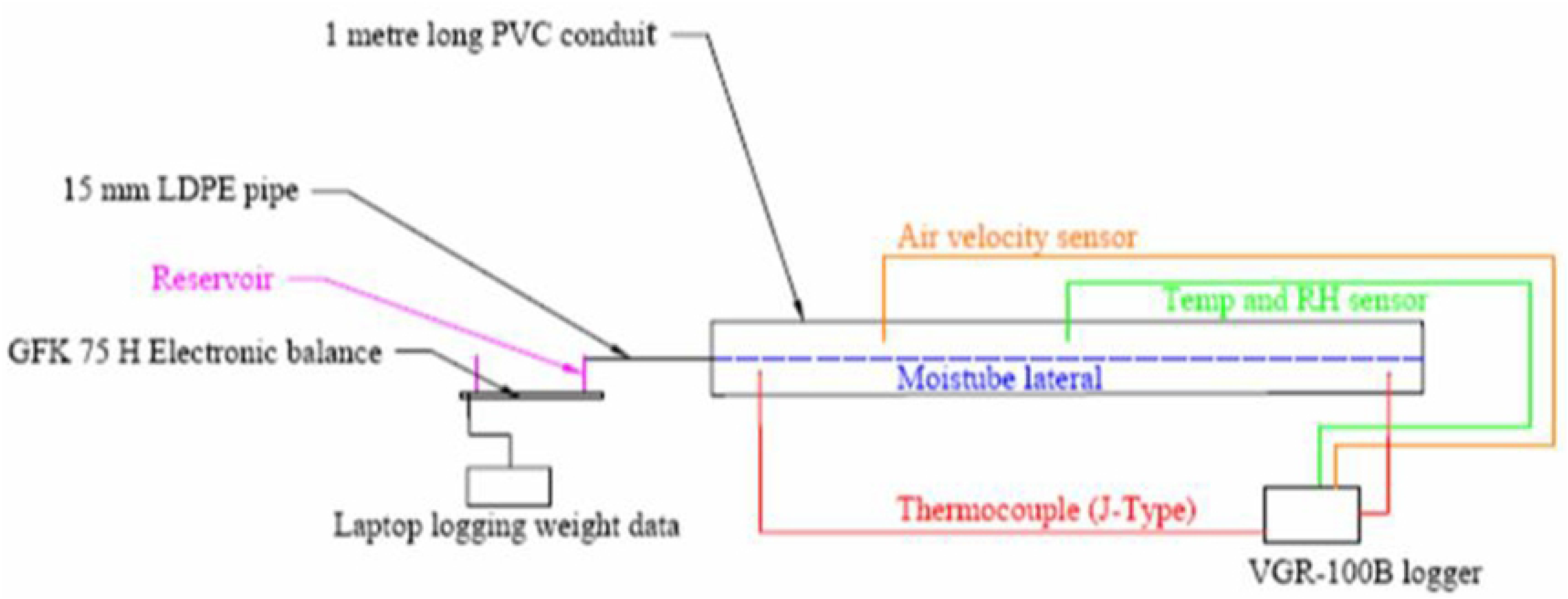
Experimental set-up.

#### 2.2.2 Data collection equipment and set-up

The equipment was assembled as shown in Figure 3. Air was blown axially to the suspended 1 m long Moistube™ lateral tubing in the PVC conduit. A flow of hot air was provided by a three speed 1800 W hot air blower. The mass flow (*m*) was measured using a GFK 75H electronic balance with a resolution of 0.001 kg. The water level in the reservoir was kept constant and at the same elevation as the Moistube lateral to eliminate the effect of water pressure head on discharge. The relative humidity (RH) was measured using the HCT01-00D sensor (E + E ELEKTRONIC ™) with a 5 - 95% RH working range, resolution of ± 2.5% RH, 2% RH accuracy, and a temperature dependency of ± 0.03% RH/°C. The temperature was measured using thermocouples (J-type) and a Pt1000 sensor (E + E ELEKTRONIC ™) with a resolution of ± 0.3°C and an accuracy of 0.1°C. Air velocity was measured using a hot film anemometer (EE 65 Series) with a working range of 0 m.s^−1^ – 20 m.s^−1^ and a resolution and accuracy of ± 0.2 m.s^−1^. The sensors were connected to a five terminal unit and 12 channel VGR-B100 (RKC Instrument ™) data logger. The logger was programmed to record average data every five-minute interval.

### 2.3 ET Model Selection

The following *ET* models were assessed as proxies for determining *E*_*d*_: (i) Morton areal, (ii) Morton point, (iii) Penman-Monteith, (iv) Priestley–Taylor, and (v) Thornthwaite (Table 1). These models are considered as universal standards for estimating *ET* (<u>9</u>). Model selection was based on the ability to accommodate all measured parameters. The Penman-Monteith model proved satisfactory since the input data variables were obtained from the experimental data comprising *T*_*a*_, (*e*_*a*_ ‒ *e*), *R*_*n*_, and *u*_*a*_.

**Table 1.** *E*_*d*_ formulations and the respective variables applied in the model(s)

| Formulation | Variables | Reference |
| --- | --- | --- |
| Morton areal | $T_a, (e_a - e), , R_n$ | (7, 10) |
| Morton point | $T_a, (e_a - e), , R_n$ | (7, 10) |
| Penman-Monteith | $T_a, (e_a - e), , R_n, u_a$ | Penman (11) cited by Donohue, McVicar (7) |
| Priestley–Taylor | $T_a, R_n$ | Priestley and Taylor (12) cited by Donohue, McVicar (7) |
| Thornthwaite | $T_a$ | Thornthwaite (13) cited by Donohue, McVicar (7) |

#### 2.3.1 Penman-Monteith model application to compute *E*_*d*_

The standardized ASCE Penman-Monteith model was used to compute *E*_*d*_. Standardized formulation aided in retaining data accuracy yet simplifying applicability. The model is defined by Equation 5 (11, 14–16):

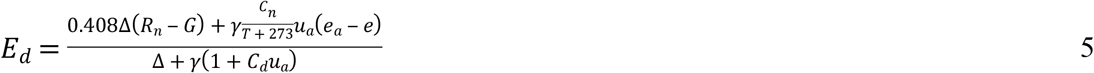

where: Δ = slope of the saturation vapour-pressure curve at air temperature (kPa.°C), *R*_*n*_ = net radiation (W.m^−2^), *e*_*a*_ ‒ *e* = vapour-pressure deficit (mbars), *e* = actual vapour pressure of air (mbars), *γ* = psychometric constant of proportionality (kPa.°C^−1^), *T* = hourly air temperature (°C), *C*_*n*_ = numerator constant for reference type and calculation time step (*C*_*n*_ = 900), and *C*_*d*_ = denominator constant for reference type and calculation time (*C*_*d*_ = 0.34) (<u>15</u>).

The energy balance equation was used to determine the evaporation energy. The *R*_*n*_ determined by Equation 6 and it was recorded in five-minute intervals:

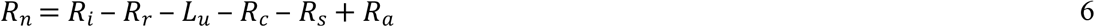

where: *R*_*i*_ = short wave solar radiation, *R*_*r*_ = reflected part of the solar energy, *R*_*c*_ is the conduction energy in air, *R*_*s*_ = incremental stored energy in the conduit, and *R*_*a*_ = advective energy. *R*_*i*_, *R*_*c*_, *R*_*s*_ and *R*_*a*_ are measured in flux units (W.m^−2^). Parameters *R*_*i*_,*R*_*r*_, *L*_*u*_, and *R*_*c*_ were neglected because the experiment was conducted in the laboratory. According to <u>Cross (17)</u> when *R*_*s*_ and *R*_*a*_ are measured in five-minute intervals which constitute “short intervals” the two parameters can be neglected since they are negligible. Furthermore, considering that the experiment was not a closed air system, it therefore invalidated the relevance of *R*_*s*_. In addition the parameter *R*_*s*_ is dependent on duration of sunshine hours (DS) and maximum possible sunshine hours available whilst the parameter *R*_*a*_ utilises the *d*_*r*_ function which is the distance between the earth and the sun (<u>18</u>), thus providing further evidence in their exclusion in Equation 7. Therefore Thus *R*_*n*_ was computed as per Equation 7:

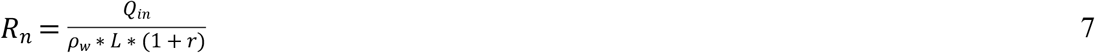

where *Q*_in_ = energy from the heater (J) (Equation 8), *p*_*w*_= density of water (kg.m^−3^), *L* = latent heat of evaporation (J.kg^−1^), and *r* = Bowen ratio (Equation 9):

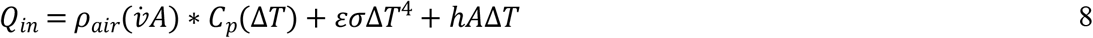

where *A* = area of the blower duct (m^2^), 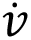 = air volumetric flow rate (m^3^.s^−1^) (product of the cross sectional area (*A*) of hot air blower duct and the average flow velocity *u*_*i*_), *C*_*p*_ = specific heat capacity of air (J.Kg^−1^. °C^−1^), *ε* is the emissivity coefficient of the PVC conduit (0.92) (19), *σ* = Stefan-Boltzmann Constant (5.670367*10^−8^ W.m^−2^.°C^−4^), ℎ = convective heat transfer coefficient (W.m^−2^.°C^−1^), *ρ*_air_ = density of air (kg.m^−3^), and Δ*T* = change in temperature along the PVC conduit heating surface (°C). The Bowen ratio is determined as follows:

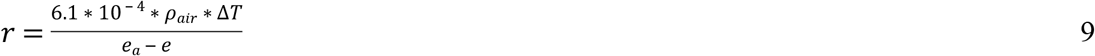

### 2.4 Statistical Analyses

A normality test was done on the three discharge data sets (*q*_*t*_) obtained from the respective *E*_*d*_ experiments using the Shapiro-Wilk normality test followed by a non-parametric Kruskal-Wallis one-way ANOVA test. All statistical analyses were done using R Studio© (<u>20</u>). A linear regression of observed and simulated values was plotted and tested for correlation using the *R*^2^ value.

## 3 Results and Discussion

### 3.1 Air Velocity (*u*_*a*_) and Evaporative Demand (*E*_*d*_)

The *E*_*d*_ values for each air velocity are presented in Table 2. The Bowen ratios (*r*) were significantly low, which means the system had a sufficient water supply. This concurs with <u>Hobbins and Huntington (5)</u> and <u>Cross (17)</u> who posited that a Bowen ratio (*r*) of less than one signifies an unlimiting hydrological supply i.e., the MTI, which is a non-water surface was relatively wet and had ample moisture to evaporate.

**Table 2.**
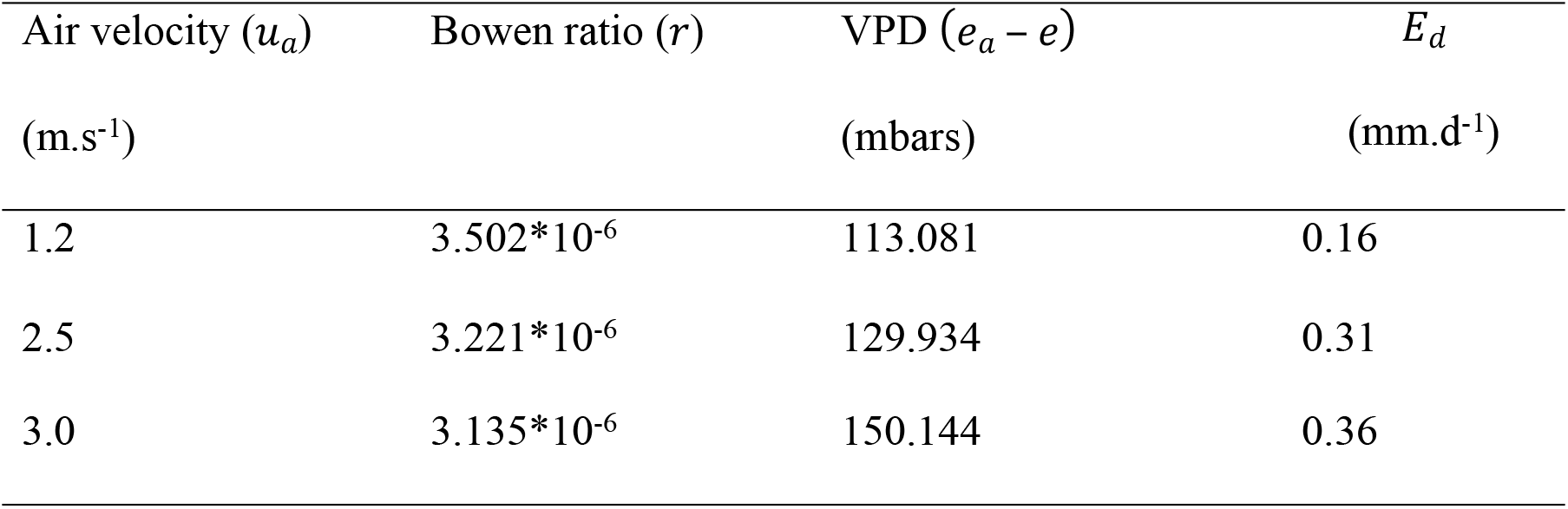
Air velocity (*u*_*a*_) measurements and the corresponding evaporative demand (*E*_*d*_)

The results (Table 2) revealed a positive correlation amongst variables *u*_*a*_, *r*, VPD and *E*_*d*_. A high air velocity (*u*_*a*_) effected a high VPD, which subsequently induced a high *E*_*d*_. The observation concurs with <u>Donohue, McVicar (7)</u> whose study attributed high evaporation rates to increased air temperatures.

### 3.2 Evaporative Demand (*E*_*d*_) and Discharge (*q*_*t*_) Relationship

The normality test (Table 3) was carried out on the evaporative demand results (*E*_*d*_ = 0.16 *mm*.*d* ^‒ 1^, 0.31 *mm*.*d* ^‒ 1^ *and* 0.36*mm*.*d* ^‒ 1^) and the respective discharge data. The sample data revealed high data skewness (*p* < 0.05). The statistical analysis revealed that there was no significant difference among the means across the three *E*_*d*_ categories (*p* = 0.05). Moistube™ discharge reaches a constant beyond a certain threshold time under the imposed negative pressure.

**Table 3.**
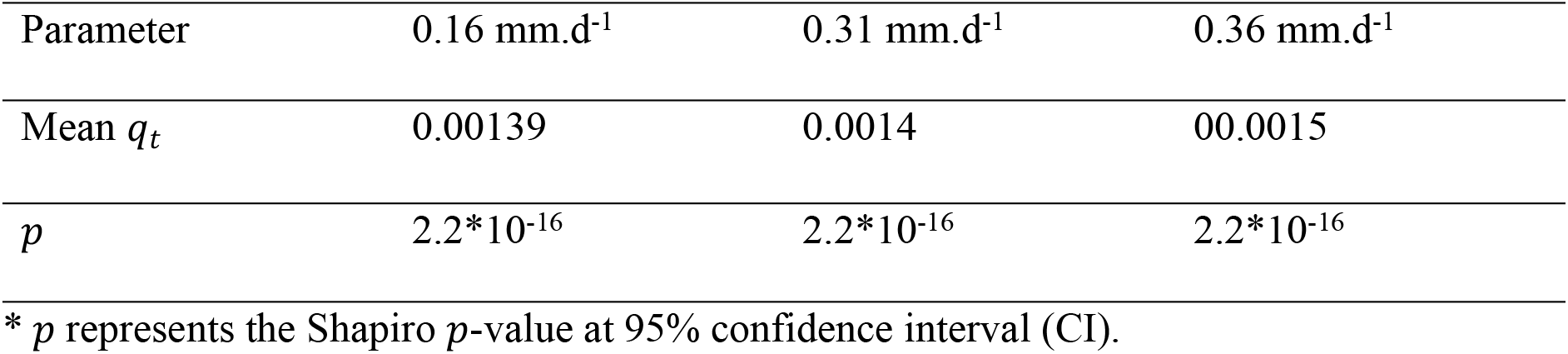
Summarised descriptive statistics for the induced *E*_*d*_ and the observed *q*_*t*_

The effect of the negative pressure was determined by establishing a relationship between mass flow rates (*m*) against the recorded *E*_*d*_. The MTI discharge under variable *E*_*d*_ was characterized by a power function (*R*^2^ = 0.62) as shown in Figure 4 over selected time scales. The overall average *q*_*t*_ vs *E*_*d*_ and modelled as in Equation 10. The equation represented average discharge (*q*_*ave*_) value for each respective *E*_*d*_ sessions, i.e., the *E*_*d*_ = 0.16 mm.d^−1^ had 1908 data points, whilst *E*_*d*_ = 0.31 mm.d^−1^ had 1608 data points, and *E*_*d*_ = 0.36 mm.d^−1^ had 120 data points. For comparative analysis purposes a normalized time scale (0 ≤ *t* * ≤ 10) was used because the last *E*_*d*_ = 0.36 mm.d^−1^ experimental conditions exceeded realistic temperature scenarios that a buried Moistube lateral can experience.

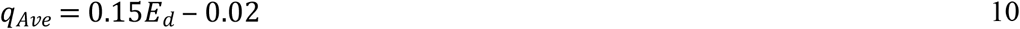

where: *q*_*Ave*_ = average discharge (l.h^−1^.m^−1^) across the respective *E*_*d*_ sessions. A linear increase in *q*_*Ave*_ relationship is observed as *E*_*d*_ increases. <u>Abu-Zreig, Abe (21)</u> in their study with ceramic pitcher pots recorded a linear discharge-evaporative demand relationship (*R*^2^ = 0.98). MTI functionality closely resembles that of ceramic pitcher pots. Equation 10 characterizes laminar flow through a porous media (MTI) and it is a function of membrane surface area (*A*), flow path length (*L*), flow duration (irrigation interval) (*t*) (<u>22</u>, <u>23</u>), and MTI hydraulic properties such as effective porosity of MTI lateral (*φ*), and inertial coefficient of MTI (*δ*) (<u>24</u>). The functional relationship is characterised by Equation 11. Equation 10 shows that at *E*_*d*_ = 0, MTI experiences zero discharge (*q*_*Ave*_ = 0), that is the MTI inertial coefficient facilitates an undisturbed fluid force.

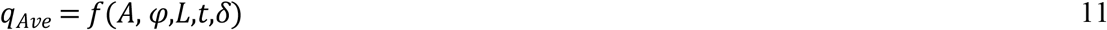

**Fig 4.**
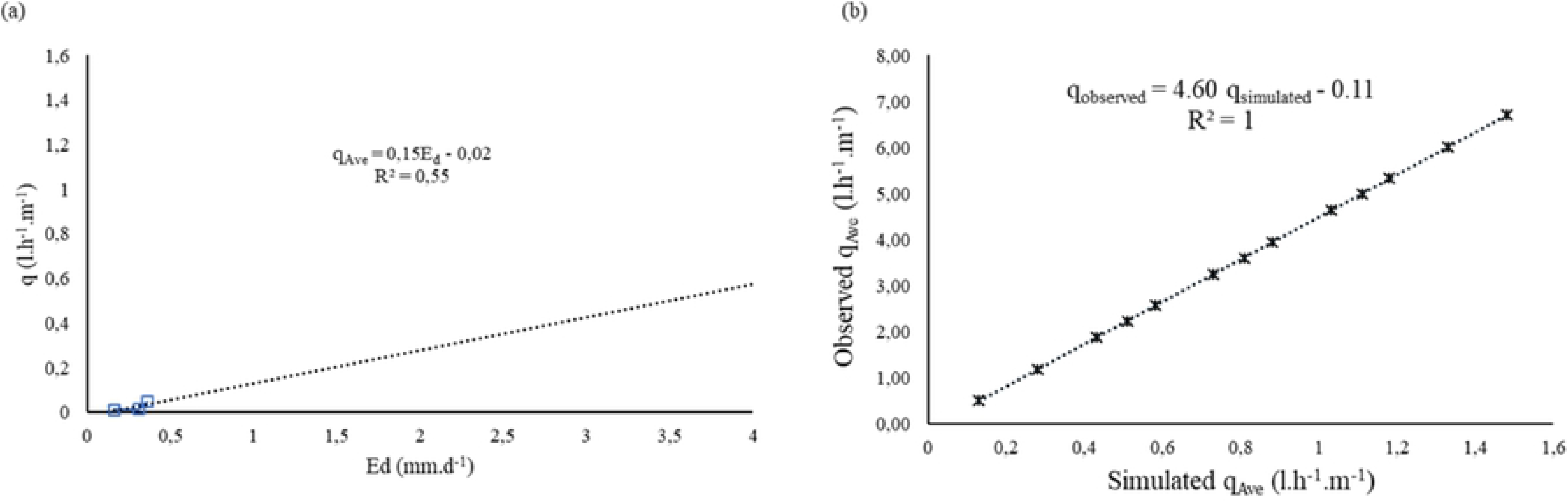
(a) Moistube™ discharge (*q*_*Ave*_) at 0 ≤ *E*_*d*_ ≤ 10 and (b) comparison between the simulated *q*_*Ave*_ vs the observed *q*_*Ave*_.

The established relationship (Equation 10) is characterized by a one sided *E*_*d*_ sensitive limit as in Equation 12. The limits were informed by the FAO evaporating power scale (<u>4</u>). The simulated *q*_*Ave*_ data had a significant difference (*p* < 0.05) across the respective regions.

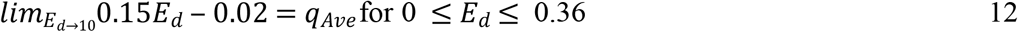

The observed (q_o_) (including the extrapolated values) and the simulated (q_sim_) (see Appendix I) were plotted on a linear regression plot and yielded a correlation value of *R*^2^ = 1 (Figure 4b). The model overestimated the *q*_*Ave*_ (PBIAS = 77 %), thus, the limits for Equation 10 applicability were defined by Equation 12.

A higher *E*_*d*_ resulted in high discharge rates. To assess the effects of *E*_*d*_ on *q* the study employed the relative discharge approach defined by Equation 13:

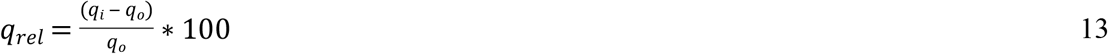

where *q*_*rel*_ = mean relative discharge (%), *q*_*i*_ = average discharge (l.h^−1^.m^−1^) at *t* * = 0 ≤ *t* * ≤ 1 h and *q*_*o*_ = average initial discharge obtained at the beginning of the experiment (l.h^−1^.m^−1^).

For *E*_*d*_ = 0.31 mm.d^−1^ there was a 10% decline in relative discharge (*q*_*rel*_) whilst *E*_*d*_ = 0.36 mm.d^−1^ recorded a 67% decline in *q*_*rel*_ over a 10 h period. These observed variations in *q*_*rel*_ between *E*_*d*_ = 0.31 mm.d^−1^ and *E*_*d*_ = 0.36 mm.d^−1^ are attributed to high *E*_*d*_ values due to increased drying power of the air in the conduit, hence effecting high discharges. <u>Yang, Tian (3)</u> postulated that in the absence of a driving pressure MTI discharge is a function of matric potential or a negative pressure. Under *E*_*d*_ = 0.16 mm.d^−1^,It was observed that over 10 h the discharge rose from 0.043 l.h^−1^.m^−1^ to 0.077 l.h^−1^.m^−1^. This effect can be attributed to a slow and gradual increase in VPD (*e*_*a*_ ‒ *e*) within the PVC conduit that effected continuous and gradual discharges. A near saturation scenario was observed from *t* = 7.5 h to *t* = 10 h under *E*_*d*_ = 0.16 mm.d^−1^ which signified a protracted equilibrium scenario whereby *e*_*a*_ ≥ *e*. For a buried Moistube™ lateral, continuous discharge is observed until water potential between soil-moisture and the water inside the MTI equilibrates. For applicability, to ensure continuous discharge beyond equilibrium points a net positive pressure is required to effect discharge as stated in studies by <u>Niu, Lü (25) and Kanda, Niu (1)</u>.

### 3.3 Discharge (*q*) vs Time (*t*) relationship

The *q* vs *t* relationship was established on a normalised time-scale (Figure 5). Statistical analysis revealed a significant difference (*p* < 0.05) in *q* over different *E*_*d*_ scenarios. The normalised run-times for each experiment are shown in Table 4. The study established a functional relationship between *q* and *t* characterized by Equation 13.

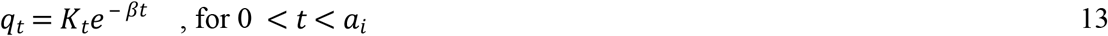

where: *q*_*t*_ = time dependent discharge, *K*_*t*_= constant of proportionality dependent on MTI hydraulic properties, MTI surface area (*A*) flow path length (*L*), and (<u>26</u>), *β* = discharge exponent, *t* * = normalised time for a specific induced *E*_*d*_, and *a*_*i*_ = normalised upper limit time range for each subsequent *E*_*d*_, value. The relationship revealed that *q* is time sensitive i.e. if MTI is continuously exposed to an imposed *E*_*d*_, MTI discharges in an exponentially decreasing trend to a point of stability at each respective normalised time.

**Table 4.**
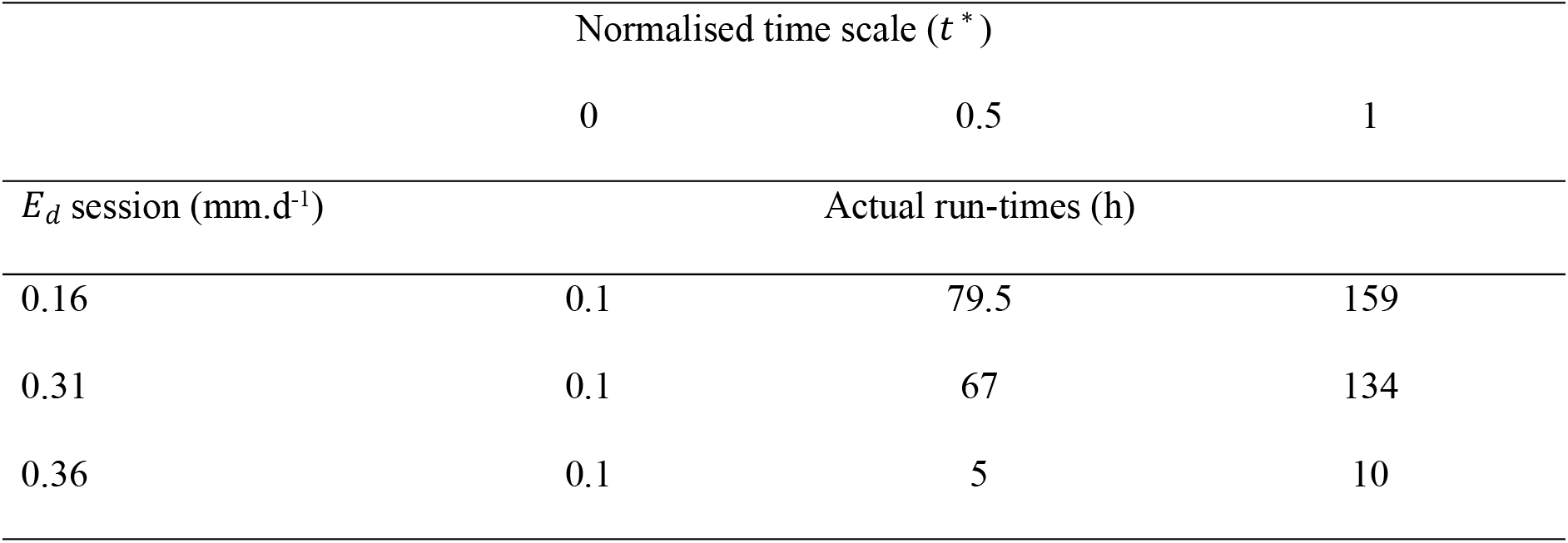
Normalised time scale (t*) vs E_d_ for the *q* vs *t* plot

**Fig 5.**
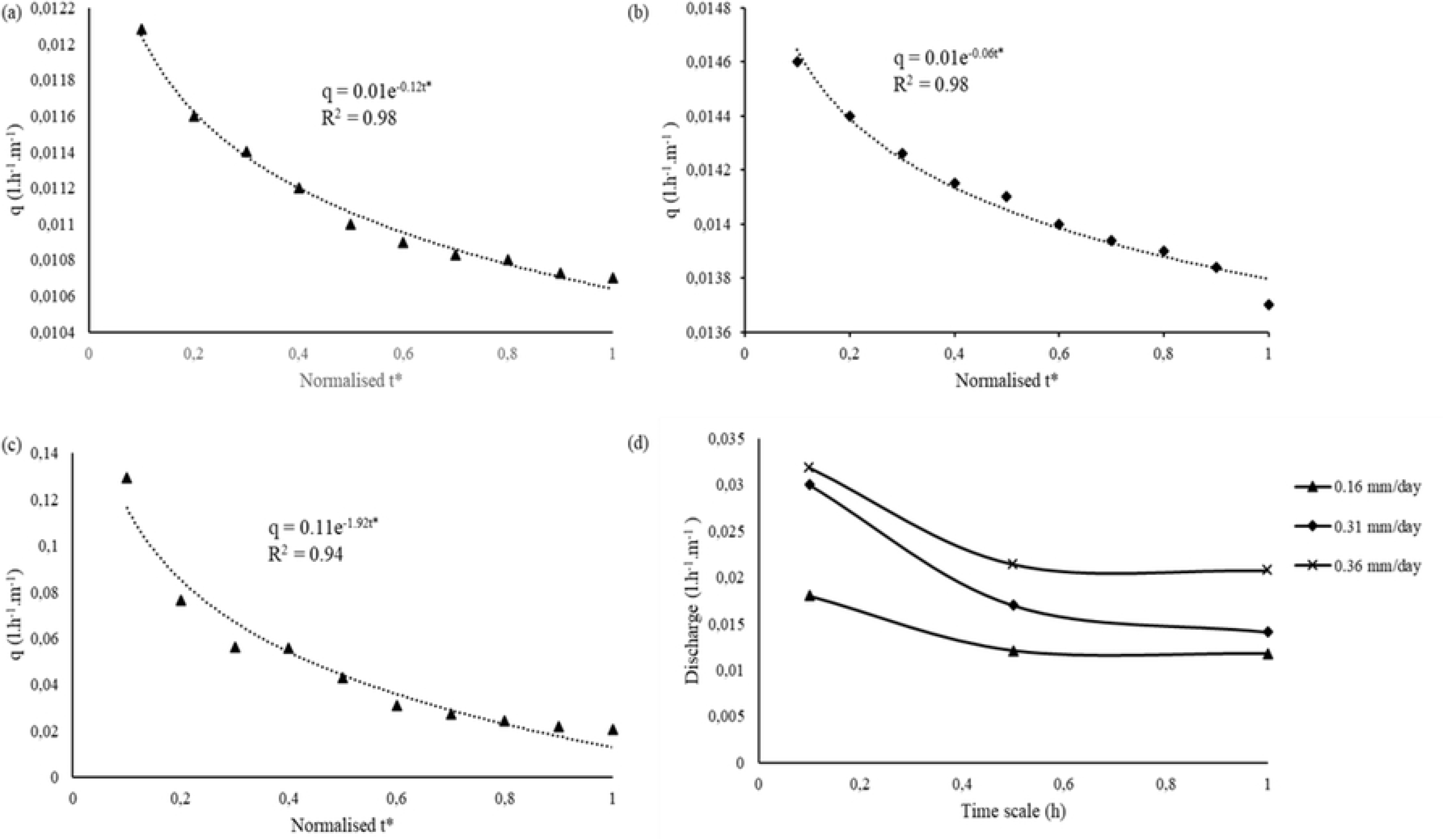
Discharge vs. normalised time (*t* *) relationship at (a) *E*_*d*_ = 0.16 mm.d-1, (b) *E*_*d*_ = 0.31 mm.d^−1^, (c) *E*_*d*_ = 0.36 mm.d^−1^, and (d) combined *q* vs *t* * plots.

For *E*_*d*_ = 0.36 mm.d^−1^ there was a steady decline in discharge. The recorded flow rate variations were approximately 0.129 l.h^−1^.m^−1^ from *t* * = 0.1 to *q* = 0.021 l.h^−1^.m^−1^ at *t* * = 10. Whereas for *E*_*d*_ = 0.16 mm.day^−1^ and *E*_*d*_ = 0.31 mm.d^−1^ discharge reached a stable steady state ranging from *q* = 0.011 l.h^−1^.m^−1^ to 0.014 l.h^−1^.m^−1^ at *t* * = 0.3 and *t* * = 0.4 respectively. These low discharge variations mimic a buried Moistube™ lateral experiencing minimal to zero discharge at low soil-water potential. <u>Niu, Zhang (27)</u> posited that at zero driving head a buried Moistube lateral reaches a stable steady state discharge after 48 hours.

The discharge for *E*_*d*_= 0.16 mm.d^−1^ at 10% of the actual time (*q* = 1.2*10^−2^) compared well with the discharge for *E*_*d*_= 0.31 mm.d^−1^ at 95 % of the actual time (*q* = 1.3*10^−2^). Flow rates reached steady state at *t* * = 0.9 for both *E*_*d*_= 0.16 mm.d^−1^ and *E*_*d*_= 0.31 mm.d^−1^. The stable flow rates describe a near saturation phenomenon whereby the air contained within the PVC enclosure reached saturation point, thus the suction effect of the imposed negative pressure had little effect on the Moistube™ discharge since the observed discharges were lower than the nominal MTI discharge of 0.3 l.h^−1^.m^−1^. <u>Yang, Tian (3)</u> asserted that discharge from a buried Moistube™ lateral reduces or stops when the matric potential of the surrounding soil approaches saturation i.e. when *ψ*_*water*_ ≤ *ψ*_*soil*_ less seepage from the Moistube™ tubing is anticipated and when *ψ*_*water*_ = *ψ*_*soil*_ there is zero seepage. Once MTI is in equilibrium with its surrounding discharge stops and only a positive driving pressure will induce discharge. A similar observation was made by (<u>21</u>).

For *E*_*d*_ = 0.16 mm.d^−1^ it was observed that there was uniform and steady decline in discharge rates from *t* =0.1 hrs to *t* * = 0.5 (33% decrease in relative discharge rate), there-after there was a steady state discharge rate (*t* * = 0.5 vs *t* * = 1) where the discharge varied from 0.0121 l.h^−1^.m^−1^ to 0.0117 l.h^−1^.m^−1^ resulting in a *q*_*rel*_ = 2.5%. The phenomenon can potentially be attributed to a stagnating VPD (*e*_*a*_ ‒ *e*) over time. The steady state discharge rates were also observed on the *E*_*d*_ = 0.31 mm.d^−1^ after *t* * = 0.5. This phenomenon indicates a situation where VPD stabilizes such that evaporation occurs over an extensively wet MTI tubing i.e. wet environment evaporation (*E*_*w*_) (<u>5</u>). For a buried MTI tubing the slow stable irrigation water release would occur in a near saturation MTI – *ψ*_*soil*_ continuum. From Figure 5, the *E*_*d*_ = 0.36 mm.d^−1^ discharge decreased rapidly due to the high drying power of the air which increased the VPD (*e*_*a*_ ‒ *e*) resulting in a high *E*_*d*_ that induces discharge. The findings concur with <u>Abu-Zreig, Zraiqat (8)</u> experiment with pitcher pots under constant head. The seepage rate of both buried pitcher pots and those in a controlled environment had a steady decrease as soil water increase. All three plots plateaued signifying a saturated micro-environment within the PVC conduit and consequently a decreased air suction effect.

## 4 Conclusions and Recommendations

Moistube performs according to the conceptual design, where it releases moisture at zero positive pressure head. At various evaporative demand scenarios the Moistube released moisture at diminishing rates. The study rejected the null hypothesis that the artificially imposed E_d_ does not cause MTI tubing discharge. The discharge rates showed an exponential decrease over operation time.

The study was an open-air experiment; hence it is recommended that the study be carried on an actual buried MTI tubing wherein *ψ* are present and influence soil-moisture movement. The *E*_*d*_ vs *q*_*Ave*_ relationship should be experimentally explored to define the actual limits of the MTI operation and *q* variations for high evaporating (ET_o_) power of the atmosphere.

When Moistube™ tubing is exposed to an evaporative demand a negative pressure develops that induces discharge. The phenomenon facilitates slow release of water thus allowing MTI tubing to be utilized for optimal field water use efficiency (fWUE), i.e., slow releasing moisture as per crop water requirements. MTI capability to maintain soil water at 80 – 90 % field capacity (FC) levels is instrumental in availing adequate crop water without having to fully saturate the soil (28). This is reflected in the study wherein discharge plateaus (constant) under *E*_*d*_ = 0.16 mm.d^−1^ and *E*_*d*_ = 0.31 mm.d^−1^. The implication highlighted was that MTI can supply water without applying a driving pressure along an irrigation line. In addition, the negative pressure system can supply water in minute quantities just below crop water requirements.

The zero-pressure head discharge phenomenon saves on irrigation energy that irrigators otherwise would incur when using other irrigation systems. The study mimicked a buried MTI tubing, wherein the VPD represented the soil matric potential. The approach can be used to model irrigation schedules based on soil matric potential (*ψ*), which subsequently avails irrigation water as per crop water requirements (CWRs), thus improving water use efficiency (WUE). Climate change destabilizes planned crop water use patterns thus, MTI tubing can be adopted in arid and semi-arid regions for intermittent water application. The *ψ* concept as investigated by the study in the form of an evaporative demand (*E*_*d*_) can be utilized by MTI tubing to facilitate water saving by timeously controlling discharge rates and thus availing water to crops when the soil-moisture drops beyond a critical level. This helps lower the yield penalties on crops that are sensitive to water deficit.

## 7 Supporting Information

This is S1 Fig 1 Conceptual active fluxes acting on a MTI lateral buried in the soil.

This is S2 Fig 2 The recorded evaporative demand experimental parameters

This is S3 Fig 3 Experimental set-up.

This is S4 Fig 4 (a) Moistube™ discharge (*q*_*Ave*_) at 0 ≤ *E*_*d*_ ≤ 10 and (b) comparison between the simulated *q*_*Ave*_ vs the observed *q*_*Ave*_.

This is S5 Fig 5 Discharge vs. normalised time (*t* *) relationship at (a) *E*_*d*_ = 0.16 mm.d^−1^, (b) *E*_*d*_ = 0.31 mm.d^−1^, (c) *E*_*d*_ = 0.36 mm.d^−1^, and (d) combined *q* vs *t* * plots.

## APPENDIX I

Data for the computed *q*_*Ave*_ values using Equation 10

| Region | Evaporating power (ET<sub>o</sub>) (14) | Computed $q_{Ave}$ |
| --- | --- | --- |
| **Humid** | 1 mm.d<sup>-1</sup> | 0.13 |
|  | 2 mm.d<sup>-1</sup> | 0.28 |
|  | 3 mm.d<sup>-1</sup> | 0.43 |
| **Sub-Humid** | 3.5 mm.d<sup>-1</sup> | 0.51 |
|  | 4 mm.d<sup>-1</sup> | 0.58 |
|  | 5 mm.d<sup>-1</sup> | 0.73 |
| **Semi-Arid** | 5.5 mm.d<sup>-1</sup> | 0.81 |
|  | 6 mm.d<sup>-1</sup> | 0.88 |
|  | 7 mm.d<sup>-1</sup> | 1.03 |
| **Arid (> 7 mm.d<sup>-1</sup>)** | 7.5 mm.d<sup>-1</sup> | 1.11 |
|  | 8 mm.d<sup>-1</sup> | 1.18 |
|  | 9 mm.d<sup>-1</sup> | 1.33 |
|  | 10 mm.d<sup>-1</sup> | 1.48 |

